# Normalized Semi-Covariance Co-Efficiency Analysis of Spike Proteins from SARS-CoV-2 variant Omicron and Other Coronaviruses for their Infectivity and Virulence

**DOI:** 10.1101/2022.11.07.515557

**Authors:** Tong Xu, Shanyue Zhou, Jun Steed Huang, Wandong Zhang

## Abstract

Spectrum-based Mass-Charge modeling is increasingly used in biological analysis. To explain statistical phenomenon with positive and negative fluctuations of amino acid charges in spike protein sequences from Omicron and other coronaviruses, we propose calculation-based Mass-Charge modeling, a normalized derivation algorithm with exact Excel and MATLAB tool involving separate quadrant extension to normalized covariance, which is still compatible with Pearson covariance co-efficiency. The number of amino acids, molecular weight, isoelectric point, amino acid composition, charged residues, mass-charge ratio, hydropathicity of the proteins were taken into consideration in the analyses, and the relative peak and dip of the average with spike protein sequences based on hydrophobic mass to isoelectric charges of amino acids were also examined. The analyses with the algorithm provide more clear insights leading to revealing underline evolving trends of the viral proteins. Spike proteins from SARS-CoV-2 variants, seasonal and murine coronaviruses were taken as representative examples in this study. The analyses demonstrate that the Mass-Charge covariance co-efficiency can distinguish subtle differences between biological properties of spike proteins and correlate well with viral infectivity and virulence.

## 1. Introduction

Spectrum-based Mass-Charge algorithms are used in analyzing real-world implementations. It comes as the trusted analytic solution but typically has a hardware implementation cost and time challenges, leading to a demand for more straightforward software calculation-based solutions using Mass-Charge theory. Mass-Charge algorithms have received significant attention in recent years and are increasingly used to solve real-world problems. Among those is a combination of two or more algorithms involving numerical algorithms, analytic calculation [1], and other computational techniques, such as artificial intelligence [2–4], gene analysis systems [5], and gene simulation [6].

Kumar and colleagues found that SARS-CoV-2 Omicron and sub-variants had a higher positive electrostatic surface potential [7]. This could increase interactions between the receptor-binding domain (RBD) of Omicron spike protein and the electro-negatively charged human angiotensin-converting enzyme 2 (hACE2). We compared Omicron spike protein and its RBD with those of Wuhan-Hu-1 (Wild type) strain. Our previous study calculated the charges of the SARS-CoV-2 spike protein sequences using the algorithm for viral infectivity and virulence [8]. In this study, the number of amino acids, molecular weight, theoretical pI, amino acid composition, charged residues, mass-charge ratio, hydropathicity of the spike proteins were all analyzed by using the improved algorithm with normalization of spike protein length.

This study is the first to use Langland’s formula to combine all the factors, including the mass, charge, isoelectric point, hydrophilic and hydrophobic properties, and equipotential of the amino acids to analyze the spike proteins from Omicron and other coronaviruses. The hydrophilic and hydrophobic properties express the ability of viral spike proteins to interact with human cells. There is a linear relationship between the isoelectric point of the amino acids and the pH value of the amino acids. The isoelectric point and pH value of the amino acid sequences change the equivalent charge potential. There is also a linear relationship between hydrophilic and hydrophobic properties of the amino acids and the heat or intrinsic energy level. The hydrophobicity and the heat/intrinsic energy change the equivalent mass. In the following sections, we use the equivalent mass and charge value as the ratio to calculate the overall Mass-Charge ratio. The viral basic reproduction numbers (R0 values) are also an essential factor for us to determine the correlation of infectivity. R0 values are the expected number of the cases directly generated by one case in a population where it assumes that all individuals are susceptible to infection [9]. R0 value is the indicator for the level of contagious infectious diseases. For example, the R0 value of smallpox is 3.5-6 (varies under different medical conditions) [10]. The R0 value for the primary strain of the SARS-CoV-2 virus is estimated to be 1.4-2.4 [11]. The detailed R0 value for each SARS-CoV-2 variant is listed in Table 1. Another statistics used in the analysis is the death rate for each variant’s virulence. A study published by the Public Health Ontario analyzed the patients infected with the Omicron variant (37296 cases) or Delta variant (24432 cases) and chose 9087 cases from each variant to analyze the death rate. Their results showed that the death rate for the Omicron variant was 0.03%, while the death rate for the Delta variant was 0.3% [12].

**Table 1:** Analysis results of spike protein from SARS-CoV-2 Wuhan strain in comparison with spike proteins from SARS-CoV-2 variants.

| Wuhan (baseline)/<br>Delta (reference) | Delta | Omicron | IHU<br>(France) | Indonesia<br>variant | Mu<br>variant | UK<br>(Alpha) | BA.5 |
| --- | --- | --- | --- | --- | --- | --- | --- |
| Positive Correlation | 99.44% | 98.09% | 98.80% | 99.59% | 99.28% | 99.58% | 98.27% |
| Negative<br>Correlation | 0.17% | 1.00% | 0.59% | 0.27% | 0.52% | 0.32% | 0.89% |
| Pearson Correlation | 0.9927 | 0.9709 | 0.9821 | 0.9932 | 0.9876 | 0.9926 | 0.9738 |
| Center<br>Convergence | 629 | 631 | 633 | 626 | 628 | 626 | 633 |
| Center Divergence | 557 | 361 | 419 | 707 | 478 | 655 | 417 |
| M/Q Span | 72 | 270 | 214 | 162 | 150 | 58 | 216 |
| Reproduction Rate | 2.08 | 7.79 | 6.17 | 4.67 | 4.33 | 1.67 | 6.23 |
| R0 Value* | 5.08 | 7.0 | 8.2 | 6.79 | 3.5 | 5.6 | 7.0 |
| Max Positive<br>Position | 518 | 518 | 516 | 516 | 517 | 516 | 518 |
| Max Negative<br>Position | 616 | 216 | 490 | 757 | 485 | 614 | 616 |
| M/Q Density | 493 | 64 | 289 | 463 | 351 | 505 | 349 |
| Virulence | 2.15% | 0.28% | 1.26% | 2.02% | 1.53% | 2.20% | 1.52% |
| Death Rate* | 0.3%-<br>3.4% | 0.06%-<br>0.3% | 0.9%-<br>2.3% | 1.0%-<br>2.3% | 2.0%-<br>3.0% | 1.3-5.3% | 0.06%-<br>0.3% |

\* The R0 value and the death rate were obtained from Wikipedia.

## 2. Normalization of Separate Quadrant Covariance

The Mass-Charge parameter is used to measure the fluctuation-term memory of time series. It relates to the autocorrelations of the time series and the derivatives of the Poincare transform of the time series or the momentum generation function at the origin [13]. Studies involving the Mass-Charge parameter were originally developed by the Fuchsian Group in solving differential equations.

Using Langland’s formula, we calculate w, h, q, and p to obtain the complete viral spike protein sequences. After the complete sequence is obtained, the N subsequences of the complete sequence are constructed with the window shift of 1, the window length of 16, and the 15 subsequences at the end to reduce the window length in turn.

In this study, we employ the Mass-Charge from Langlands program in below formula (1).

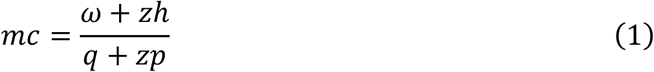

The full sequence of Mass-Charge is defined as *S* = {*mc*_1_, *mc*_2_,…, *mc_n_*), the n was the length of a full viral spike protein. In the above formula, ω is the molecular mass; *h* is the hydrophilic and hydrophobic index; q is the charge; p is the isoelectric point; z is equivalent to Finsler distance between viral spike protein and human cell receptor, typical range from 0.01 to 10000 (nm), and the default is 1. Excel is used to calculate the default, MATLAB is used to calculate the non-default distance variation map, and cross-check with the Excel calculation.

### 2.1. The Convergence and Divergence

The convergence of the viral protein are calculated by formula (2), the divergence of the viral protein are calculated by formula (3).

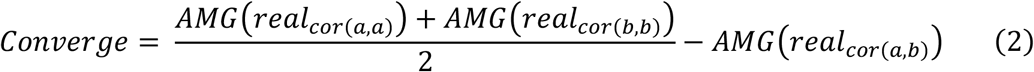

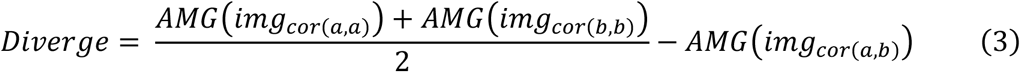

In formula (2) and formula (3), a and b are two different viral protein sequences; AMG is average moving gate function; *real*_*cor*(*a,b*)_ is the converging covariance of viral protein a and viral protein b, *real*_*cor*(*a,a*)_ and *real*_*cor*(*b,b*)_ are the special case which is that viral protein a was equal to viral protein b, they are calculated from one viral protein sequence with reference to itself instead of to other baseline one. *img*_*cor*(*a,b*)_is the diverging covariance of a and b; *img*_*cor*(*a,a*)_ and *img*_*cor*(*b,b*)_ also the special case of *img*_*cor*(*a,b*)_ We calculate from formula (4) and formula (5).

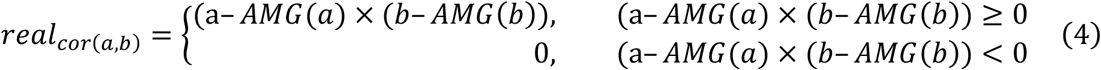

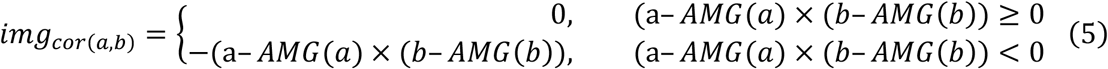

The AMG operator is defined in here, the arithmetic mean with moving window length function.

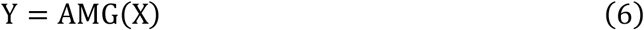

In the operator of formula (6), X = {*x_i_*|*i* ∈ [1, *n*]}, Y = *y_i_*|*i* ∈ [1, *n*]}. The calculation of AMG operator is formula (7).

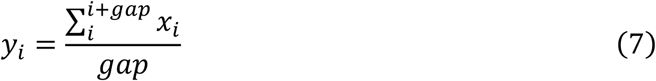

### 2.2. The Positive Correlation and Negative Correlation

Positive Correlation:

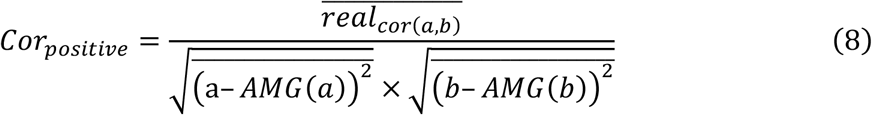

Negative Correlation:

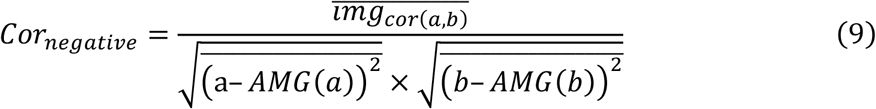

Pearson Correlation:

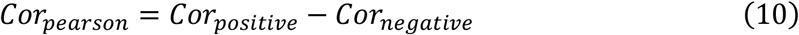

### 2.3. The Center Gravity Positive, Center Gravity Negative and M/Q Span

#### Center Gravity Positive

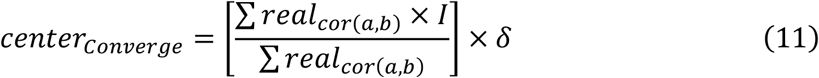

[] operator is Rounding operator; *δ* is percentage, due to different length of viral proteins;

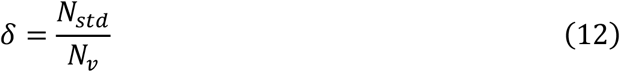

*N_std_* is the standard baseline viral protein sequence length, *N_v_* is now viral protein sequence. *I* is a sequence from 1 to the length of viral protein sequence.

#### Center Gravity Negative

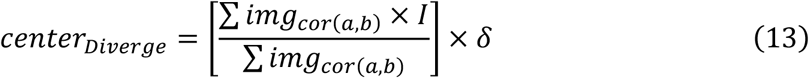

#### M/Q Span

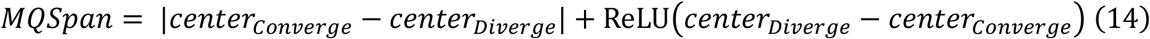

Here ReLU(x) is defined as Max (0, x).

### 2.4. Infective Reproduction Rate and Max Position

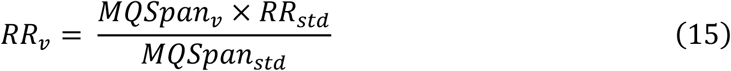

*RR_std_* is the standard baseline virus sequence infective reproduction rate; *RR_v_* is estimated viral reproduction rate; *MQSpan_std_* is the viral protein sequence standard M/Q Span; *MQSpan_v_* is estimated viral protein sequence M/Q Span. Reproduction rate here is used to approximate the observed R0 number.

Max Positive Position

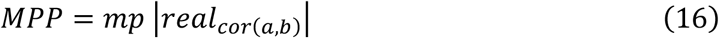

The “mp||” is the operator that get the position of max value in *real*_*cor*(*a,b*)_. Max Negative Position

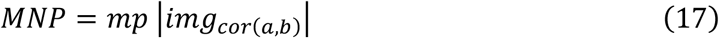

### 2.5. M/Q Density and Virulence

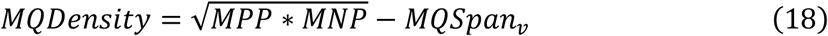

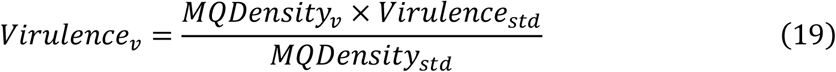

*MQDensity_std_* is the standard baseline viral protein sequence M/Q density; *MQDensity_v_* is now viral protein sequence M/Q density; *Virulence_std_* is the viral standard virulence; *Virulence_v_* is current viral virulence.

## 3. Excel calculations of normalized semi-covariance for spike proteins from SARS-CoV-2 and other coronaviruses

To prove the usage of the simplified Mass-Charge variances, we compare the correlation of SARS-CoV-2 viral spike protein with other coronaviral spike proteins [14]. Since Excel is capable of handling ReLU, we simplify the calculation with normalization between two variables only, although more than two can also be done. Since each coronaviral protein from animal or human has different electrical charge level [15], we normalize the covariance by the variance respectively (so that the comparison is focused on the pure difference) [16]. We calculated the whole sequences of spike proteins [17] and plotted the curve forward starting from the low end to high end (from 1 to 1500 or 1900 depends on the length of the spike proteins). By using the moving window of 16 neighboring amino acids [18], we calculated the covariance and average over the same area of the sequences to make the curve more visually smooth for easily comparisons [19]. Figures 1 to 3 show the calculation results for the most related spike protein sequences out of past years from murine coronaviruses, from SARS-CoV-2 and the variants Omicron, Delta, IHU, and from seasonal coronaviruses OC43, 229E, HKU1, and NL63 for human (Table 2) [20–22].

**Figure 1.**
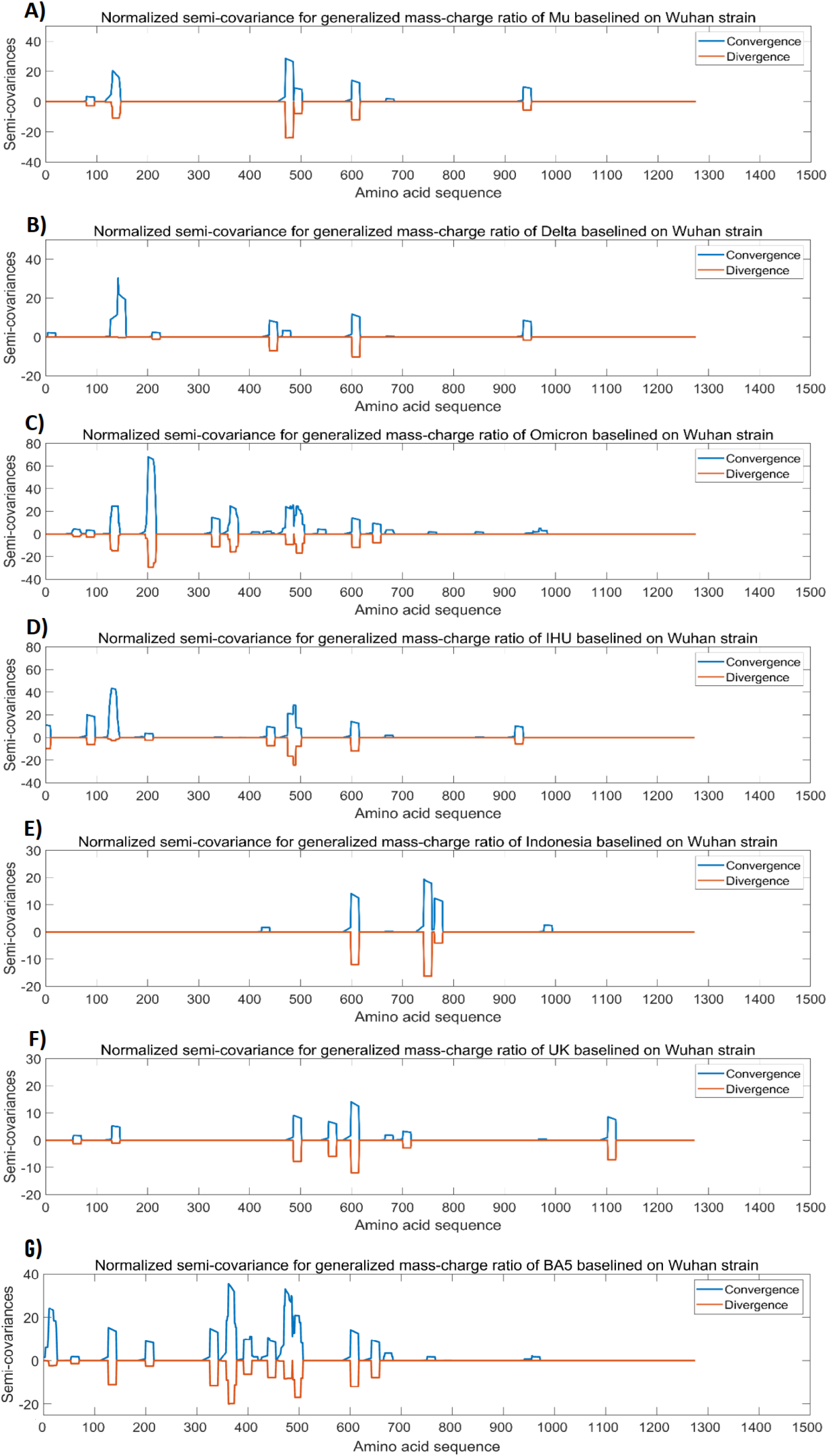
The analysis results of spike protein from Wuhan strain SARS-CoV-2 in comparison with spike proteins from SARS-CoV-2 variants. The peaks and the dips in the graphs represent variations in Mass-Charge ratio of the amino acids of the spike proteins, meaning that mutations occur in this region resulting in changes in charge/mass of the amino acids. The wider peaks or dips represent the changes involved in more amino acid changes of the spike proteins. The peaks indicate positive correlation and the dips indicate negative correlation. The dips mean opposite changes in the charge/mass of the amino acids of the spike proteins. The dips may have stronger impact on the biological function of the proteins than the peaks. Panel A: Normalized semi-covariance for generalized mass-charge ratio of Mu variant (B.1.621) spike protein baselined on Wuhan strain spike protein. Panel B: Normalized semi-covariance for generalized mass-charge ratio of Delta variant (B.1.617) spike protein baselined on Wuhan strain spike protein. Panel C: Normalized semi-covariance for generalized mass-charge ratio of Omicron variant spike protein baselined on Wuhan strain spike protein. Panel D: Normalized semi-covariance for generalized mass-charge ratio of IHU variant (B.1.640) spike protein baselined on Wuhan strain spike protein. Panel E: Normalized semi-covariance for generalized mass-charge ratio of Indonesia variant (B.1.446.2) spike protein baselined on Wuhan strain spike protein. Panel F: Normalized semi-covariance for generalized mass-charge ratio of Alpha/UK variant (B.1.1.7) spike protein baselined on Wuhan strain spike protein. Panel G: Normalized semi-covariance for generalized mass-charge ratio of BA.5 (B.1.1.529+BA) spike protein baselined on Wuhan strain spike protein.

**Figure 2.**
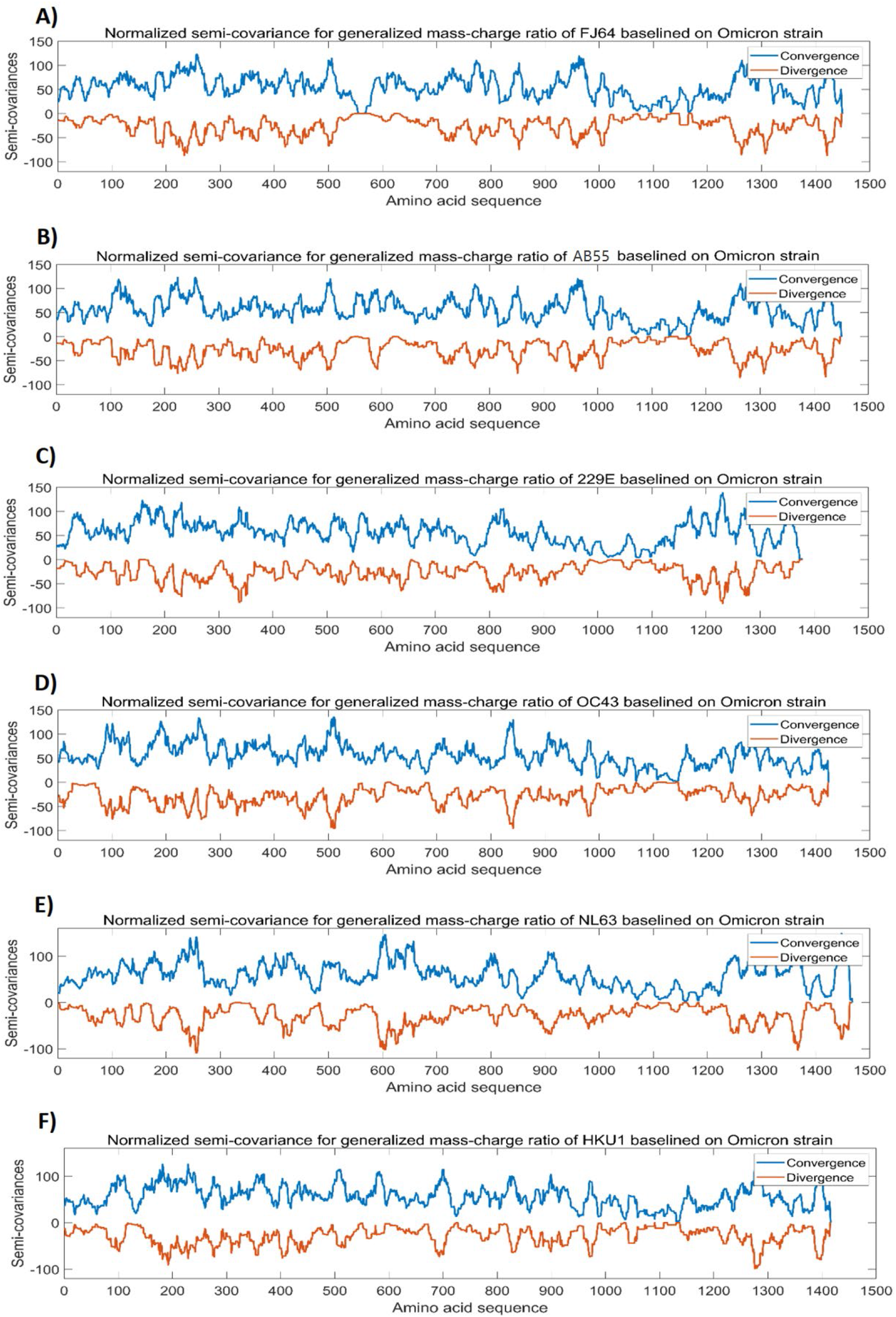
The analysis results for Omicron spike protein in comparison with spike proteins from murine coronaviruses and human common cold coronaviruses. The peaks and the dips in the graphs represent variations in Mass-Charge ratio of the amino acids of the spike proteins, meaning that mutations occur in this region resulting in changes in charge/mass of the amino acids. The wider peaks or dips represent the changes involved in more amino acid changes of the spike proteins. The peaks indicate positive correlation and the dips indicate negative correlation. The dips mean opposite changes in the charge/mass of the amino acids of the spike proteins. The dips may have stronger impact on the biological function of the proteins than the peaks. Panel A: Normalized semi-covariance for generalized mass-charge ratio of murine coronavirus FJ64 spike protein (Genbank: FJ647218.1) baselined on Omicron spike protein. Panel B: Normalized semi-covariance for generalized mass-charge ratio of murine coronaviral spike protein (GenBank: AB551247.1) baselined on Omicron spike protein. Panel C: Normalized semi-covariance for generalized mass-charge ratio of spike protein from human common cold coronavirus 229E baselined on Omicron spike protein. Panel D: Normalized semi-covariance for generalized mass-charge ratio of spike protein from human common cold coronavirus OC43 baselined on Omicron spike protein. Panel E: Normalized semi-covariance for generalized mass-charge ratio of spike protein from human common cold coronavirus NL63 baselined on Omicron spike. Panel F: Normalized semi-covariance for generalized mass-charge ratio of spike protein from human common cold coronavirus HKU1 baselined on Omicron spike protein.

**Figure 3.**
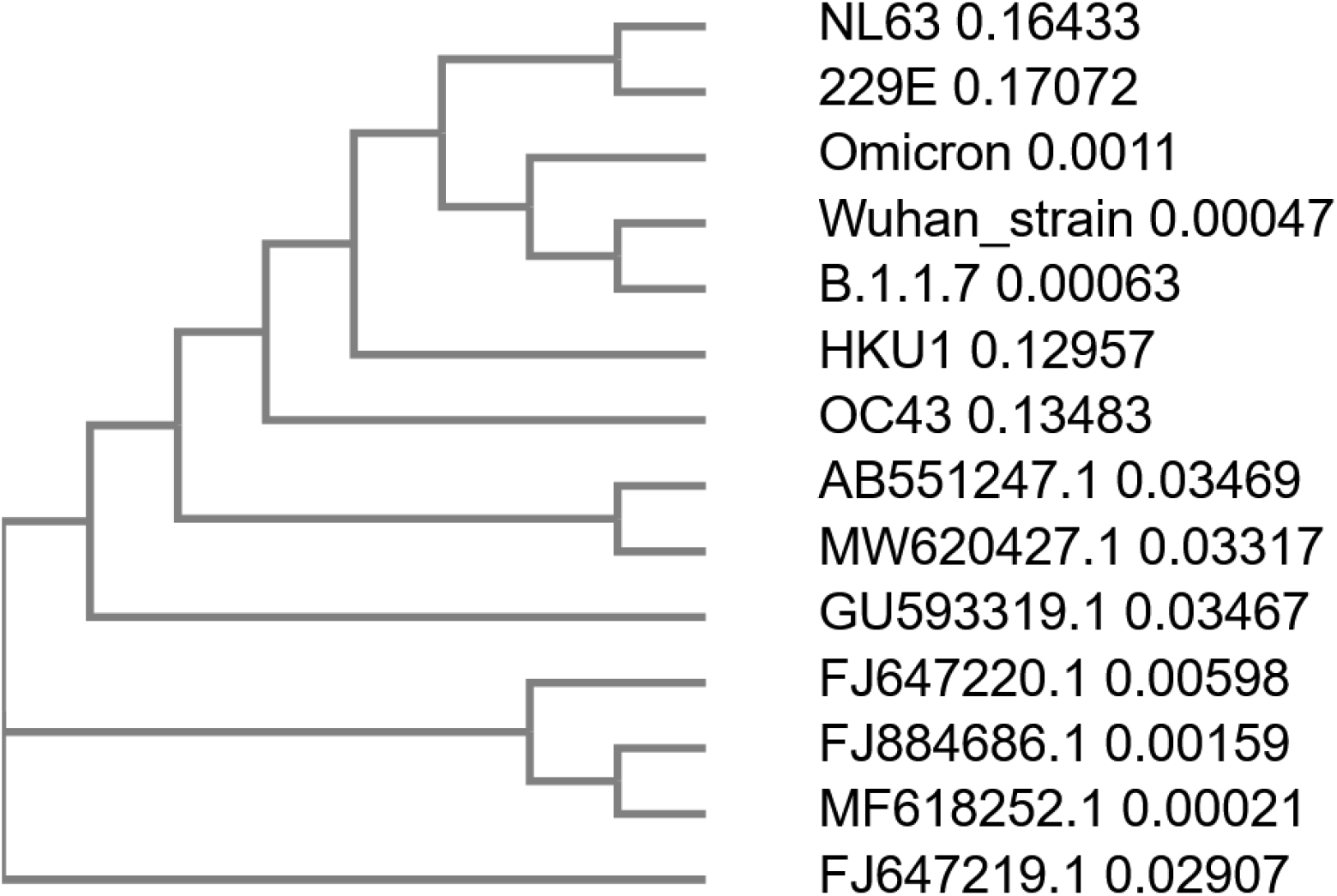
Phylogenetic tree for complete genomes of SARS-COV-2 (Wuhan strain, B.1.1.7, and Omicron), human common cold coronaviruses (NL63, 229E, OC43, and HKU1), and murine coronaviruses (Genbank Accession #: AB551247.1, MW620427.1, GU593319.1, FJ647220.1, FJ884686.1, MF618252.1 and FJ647219.1) by using Clustal Omega.

**Table 2:** Analysis results of Omicron spike protein in comparison with spike proteins from human seasonal common cold coronaviruses

| Omicron (baseline)/OC43 (reference) | OC43 | 229E | NL63 | HKU1 |
| --- | --- | --- | --- | --- |
| Year: 2021 | 1953 | 1965 | 2004 | 2004 |
| Positive Correlation | 59.06% | 56.80% | 53.97% | 59.38% |
| Negative Correlation | 20.32% | 19.93% | 21.87% | 20.46% |
| Pearson Correlation | 0.3874 | 0.3686 | 0.3210 | 0.3893 |
| Center Convergence | 695 | 686 | 681 | 674 |
| Center Divergence | 569 | 629 | 632 | 611 |
| M/Q Span | 126 | 57 | 49 | 63 |
| Reproduction Rate | 1.56 | 0.71 | 0.61 | 0.78 |
| Max Positive Position | 1180 | 1112 | 1197 | 1171 |
| Max Negative Position | 274 | 352 | 432 | 1290 |
| M/Q Density | 443 | 569 | 670 | 1166 |
| Virulence | 0.020% | 0.026% | 0.030% | 0.053% |
| Death Rate* | 0.02% | 0.03% | 0.045% | 0.10% |

\* The data were obtained from Wikipedia.

### 3.1. The analysis results for Wuhan strain spike protein in comparison with spike proteins from SARS-CoV-2 variants

We compared the spike protein of Wuhan strain SARS-CoV-2 (NC_045512.2) with the spike proteins from SARS-CoV2 variants including Mu (B. 1.621; GISAID: EPI_ISL_4029606), Delta (B.1.617; GISAID: EPI_ISL_1731198), Omicron (GISAID: EPI_ISL_6951145), IHU (B. 1.640; GISAID: EPI_ISL_8416940), Indonesia variant (B.1.466.2; GenBank: QTS26735), Alpha/UK variant (B.1.1.7; EPI_ISL_744131), and BA.5 (B.1.1.529+BA; EPI_ISL_12464782). For the comparison, Wuhan strain spike protein is used as a baseline, and Delta variant used as a reference point. We set *RR_std_* =2.08 (Delta variant), and *Virulence_std_* =2.15% (Delta variant). The analyses results are presented in Figure 1. It appears that most changes of the Mass-Charge ratios (the peaks and dips in Figure 1A to 1F) occur in the N-terminal half or the middle region of the spike proteins but rarely occurs in the C-terminal half of the spike proteins. The N-terminal half or the S1 subunit of the spike proteins carries the receptor-binding domain (RBD) of the spike protein while the middle region of the spike protein may have the function that directly or indirectly affects the interactions of RBD to the ACE2 receptor. Mutations in the surrounding region of the RBD may have epistatic effect on the binding of the spike protein to the ACE2 as we reported previously [23]. Epistasis is the combinatory effect of two or more mutations in a genome [24]. Epistatic mutations may allow spike protein to adopt a specific conformation for better or more efficient interaction of RBD with ACE2 [23]. The RBD is located between 331 and 528 amino acid residues of the spike proteins [23]. All the variants (including Figure 1A to 1F) show the changes of Mass-Charge ratios in the RBD region of the spike proteins as compared to Wuhan strain spike protein. These changes in the RBD region affect the infectivity and transmissibility of the viral variants. It is also noticed that the changes of Mass-Charge ratios frequently occur closely on both sides of the RBD region in all the variants (Fig. 1A to 1D & 1F) except the variant B.1.466.2 (Fig. 1E). Overall, there are more changes in the RBD regions of the Omicron and its subvariant BA.5 spike proteins (Fig. 1C and 1G) as compared to others (Fig. 1A, 1B, 1D, 1E, and 1F). As said above, these Mass-Charge ratio changes on both sides of RBD may have direct or indirect impacts on the interactions of RBD with ACE2 receptor. It is also noted that there are not many changes in the C-terminal region of the variant spike proteins (Fig. 1A to 1F), indicating that this region is relatively stable or conserved during evolution.

Further to the data presented in Figure 1, Table 1 shows the detailed analysis results for the comparison of Wuhan strain spike protein with the spike proteins from SARS-CoV-2 variants. In Table 1 (also in Tables 2 and 3 as well), the Center Convergence is the center point of positive momentum, meaning that the Mass-Charge ratio’s positive momentum on both sides of the spike protein sequence is equivalent; while the Center Divergence represents the center point of negative momentum, meaning that the Mass-Charge ratio’s negative momentum on both sides of the sequence is equivalent. The M/Q Span is the non-Euclidean distance between the center of convergence and the center of divergence. The Reproduction rate in the table is the calculated result from the analyses. Max Positive Position is the maximal point of positive momentum, meaning that the Mass-Charge ratio on this point reaches the maximal positive value; while the Max Negative Position is the maximal point of negative momentum, meaning that the Mass-Charge ratio on this point reaches the maximal negative value. M/Q Density is the non-Euclidean height between the center of convergence and the center of divergence. Virulence in the Tables is the calculated results from this algorithm for the viruses. From the analysis results presented in Table 1, it appears that the calculated viral reproduction rates are very close to the R0 values reported in the literature. In particular, the calculated reproduction rate for Omicron (7.79) is almost identical to the reported R0 value for Omicron (7.0) in the literature, suggesting that the calculated result by using the algorithm based on the Mass-Charge changes in the amino acids of the spike proteins may accurately predict the infectivity of the viral variant. More importantly, the virulence in Table 1 calculated by using the algorithm is closely within the range of death rates reported in the literature, indicating that the calculated virulence of the viral variants may predict the death rate caused by the viral variants. The calculated virulence of Omicron subvariant BA.5 is higher to that of Omicron in Table 1. A new study using multiscale investigations published in October 13, 2022 suggests that the risk of BA.4/5 to global health is greater than that of original BA.2 and that BA.4/5 is more pathogenic than BA.2 [25], which is consistent with our calculation. However, due to popular vaccination/boosters and advanced COVID therapies available, the actual death rate caused by BA.5 infection appears to be similar to that of Omicron (Table 1). BA.5 may have immune escape from current vaccination and therapies, but the symptoms and severity of the disease caused by BA.5 infection may be reduced by available vaccination and therapies [25].

**Table 3:**
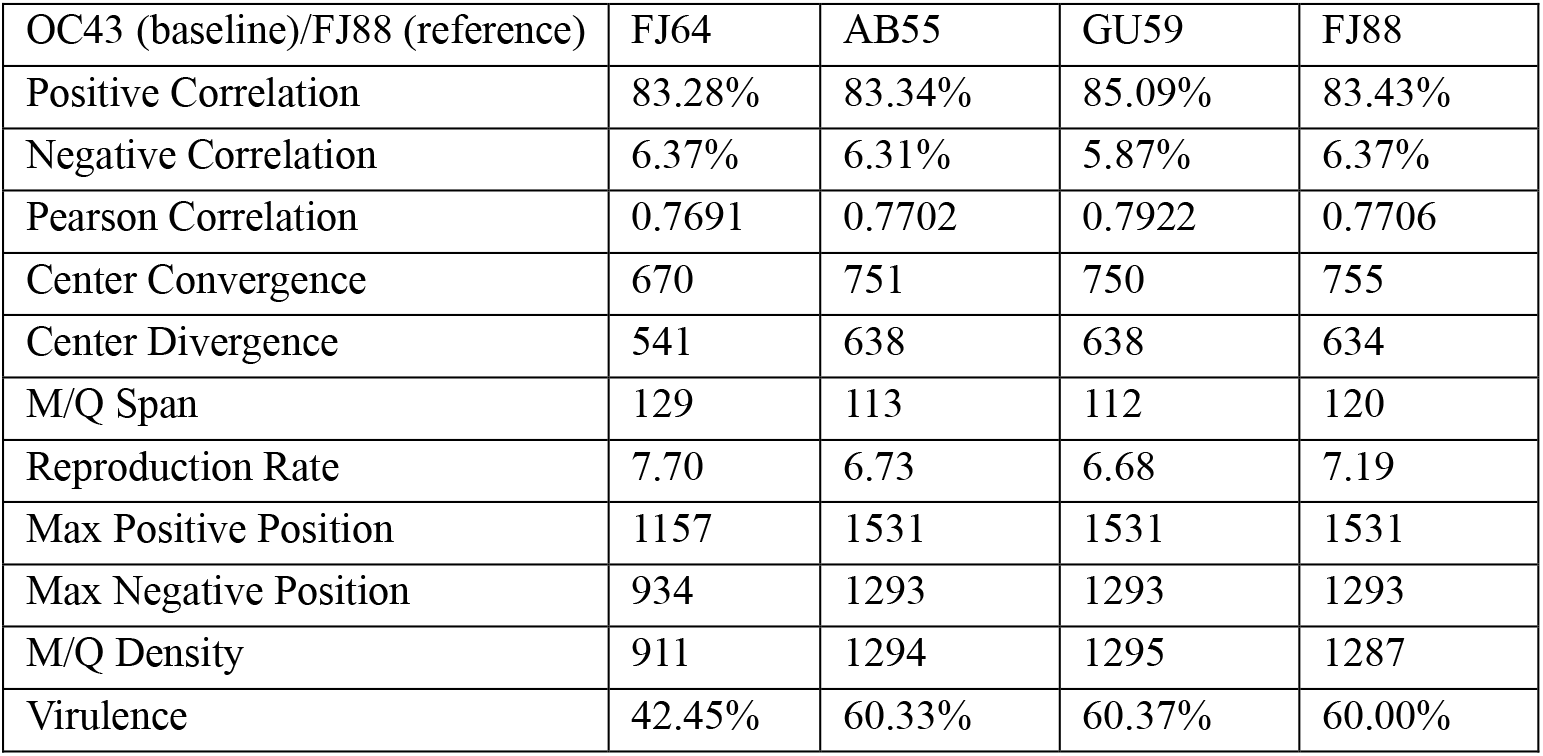
Analysis of OC43 spike protein in comparison with spike proteins from murine coronaviruses.

### 3.2. Analysis results for Omicron spike protein in comparison with spike proteins from human common cold coronaviruses and murine coronaviruses

We compared the Omicron spike protein with the spike proteins from murine coronaviruses FJ64 (Murine coronavirus RA59/R13; Genbank: FJ647218.1) and AB55 (GenBank: AB551247.1) as well as from human common cold (seasonal) coronaviruses 229E (GenBank: NC_002645.1), OC43 (GenBank: MN488635.1), NL63 (GenBank: KY554970.1), and HKU1 (GenBank: MN488637.1). For this comparison, we use Omicron spike protein as the baseline and OC43 spike protein as a reference. We set *RR_st_d* =1.56 (OC43), *Virulence_std_*=0.02% (OC43). Figure 2 shows the analysis results for Omicron spike protein in comparison with the spike proteins from murine coronaviruses and human common cold coronaviruses. Unlike the data presented in Figure 1, there are more changes in Mass-Charge ratios between Omicron spike protein and the spike proteins from the murine and seasonal coronaviruses as shown in Figure 2. Phylogenetic analysis of the viral genomes shows that Omicron and variant B.1.1.7 are closely related to Wuhan strain virus as shown in Fig. 1C and 1F; but, as compared to Wuhan strain virus and its variant B.1.1.7, Omicron is closer to seasonal coronaviruses NL63 and 229E and then to HKU1 and OC43. The phylogenetic analysis also shows that OC43 and HKU1 as well as NL63 and 229E are closer to murine coronaviruses in evolution (Fig. 3). However, these differences are not apparently observed in Fig. 2. Overall, unlike data presented in Fig. 1, there are a lot of more dissimilarities between Omicron spike protein and the spike proteins from human common cold coronaviruses and murine coronaviruses (Fig. 2), suggesting distant relationship between Omicron and these coronaviruses in evolution.

Further to the data presented in Fig. 2, Table 2 shows the detailed analysis results for the comparison of Omicron spike protein with the spike proteins from human common cold coronaviruses. As compared to the centers of convergence presented in Table 1 (626 to 633) for SARS-CoV-2 variants, the centers of convergence for the spike proteins from seasonal coronaviruses is shifted to the right (674 to 695) of the sequence. The centers of divergence for spike proteins from SARS-CoV-2 variants are scattered from 361 to 707 (Table 1); while the centers of divergence for the spike proteins from seasonal coronaviruses are more located at the region from 569 to 632 (Table 2). The max positive positions for SARS-CoV-2 variants are centralized to 516 to 518 (Table 1); while the max positive positions for seasonal coronaviruses are centralized to 1112 to 1197 (Table 2). The max negative positions for SARS-CoV-2 variants and seasonal coronaviruses are mostly scattered on the N-terminal regions of spike proteins (216 to 757) except HKU1 (1290) (Tables 1 and 2). More importantly, the virulence calculated by the algorithm using the parameters for the common cold coronaviruses is very closely to the actual death rates reported in the literature (Table 2). This further demonstrates that the calculated virulence may predicate the virulence of the viruses, i.e., the death rate caused by the coronavirus infection.

### 3.3. The analysis results for OC43 spike protein in comparison with spike proteins from murine coronaviruses and seasonal coronaviruses

We compared OC43 spike protein with the spike proteins from MW62 (murine coronavirus MHV-3; GenBank: MW620427.1), FJ64 (GenBank: FJ647218.1), AB55 (GenBank: AB551247.1), GU59 (GenBank: GU593319.1), FJ88 (GenBank: FJ884686.1), and HKU1 (GenBank: MN488637.1). In this comparison, OC43 spike protein is used as the baseline and the FJ88 spike protein is used as a reference. We set *RR_std_* =1.0 (FJ88), *Virulence_std_*=60% (FJ88). The murine coronaviruses are murine hepatitis virus (MHV) and are positive single-stranded RNA coronaviruses of ~31kb. These viruses are highly infectious and fetal. MHV infection of rodents can cause ~60% death and the mortality can reach 100% in infant mice [26,27]. The standard virulence is thus set at 60% in this analysis, but the reference for virulence can be changed upon the viruses to be used in the analysis. Figure 4 shows the analysis results of OC43 spike protein in comparison with the spike proteins from murine coronaviruses and seasonal coronavirus HKU1. It can be seen from Figure 4 that the similarity of the Mass-Charge ratios between OC43 spike protein and murine/HKU1 coronaviral spike proteins is closer as compared to those between Omicron and seasonal/murine coronaviruses presented in Fig. 2, but the similarity in Fig. 4 is less than those between Wuhan strain virus and its variants presented in Fig. 1. It is also noted that the patterns of Mass-Charge ratios displayed in Fig. 4A to 4E are highly similar to each other. Another important observation is that the variations in the C-terminal part of the spike protein sequences are smaller than the variations in the N-terminal region and the middle region of the spike protein sequences (Fig. 4), further indicating that this region is relatively stable in terms of Mass-Charge ratios and protein sequence.

**Figure 4.**
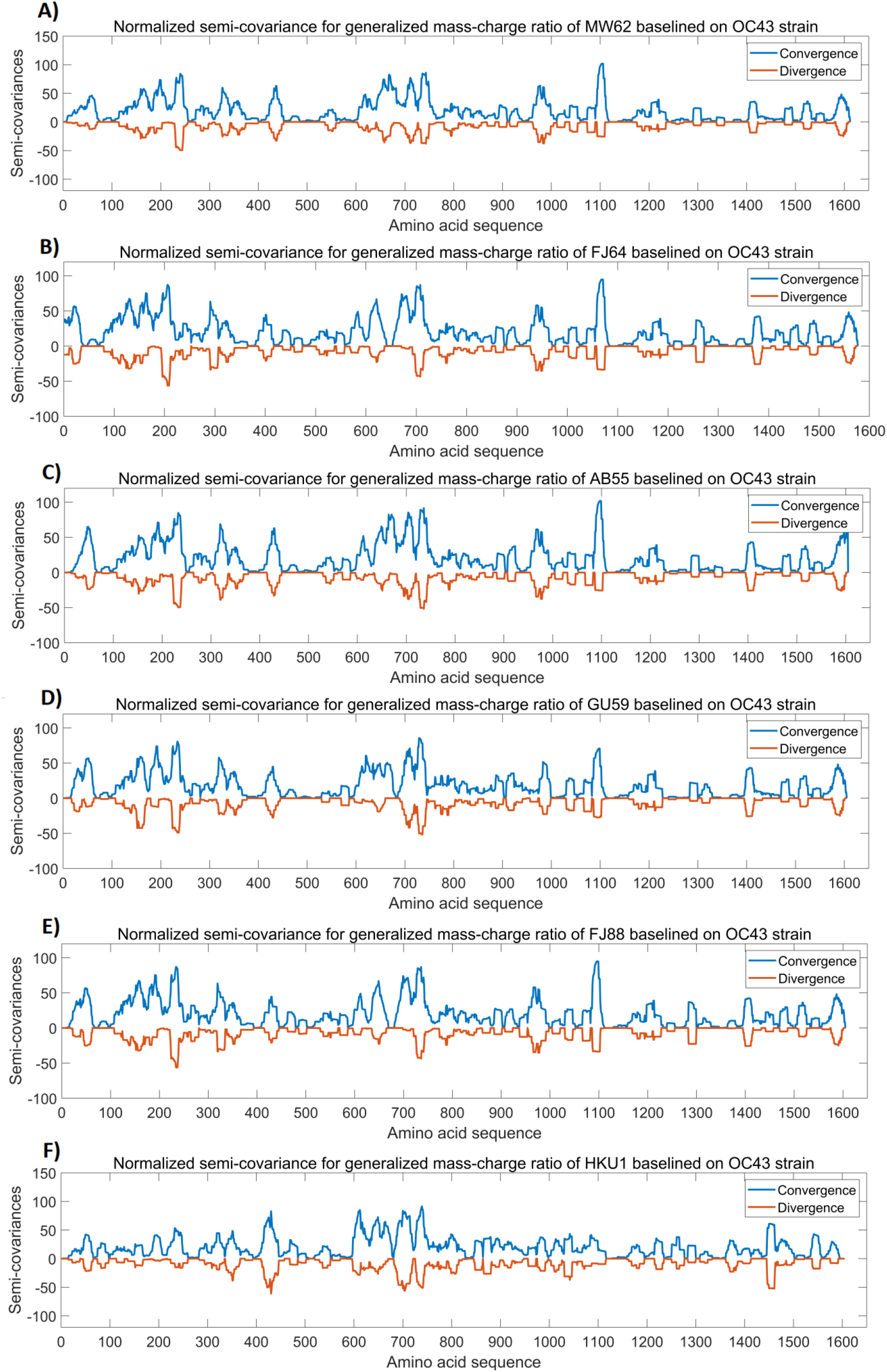
The analysis results of OC43 spike protein in comparison with spike proteins from other coronaviruses. The peaks and the dips in the graphs represent variations in Mass-Charge ratio of the amino acids of the spike proteins, meaning that mutations occur in this region resulting in changes in charge/mass of the amino acids. The wider peaks or dips represent the changes involved in more amino acid changes of the spike proteins. The peaks indicate positive correlation and the dips indicate negative correlation. The dips mean opposite changes in the charge/mass of the amino acids of the spike proteins. The dips may have stronger impact on the biological function of the proteins than the peaks. Panel A: Normalized semi-covariance for generalized mass-charge ratio of murine coronaviral spike protein (GenBank: MW620427.1) baselined on OC43 spike protein. Panel B: Normalized semi-covariance for generalized mass-charge ratio of murine coronaviral FJ64 (GenBank: FJ647218.1) spike protein baselined on OC43 spike protein. Panel C: Normalized semi-covariance for generalized mass-charge ratio of murine coronaviral AB55 (GenBank: AB551247.1) spike protein baselined on OC43 spike protein. Panel D: Normalized semi-covariance for generalized mass-charge ratio of murine coronaviral GU59 (GenBank: GU593319.1) spike protein baselined on OC43 spike protein. Panel E: Normalized semi-covariance for generalized mass-charge ratio of murine coronaviral FJ88 (GenBank: FJ884686.1) spike protein baselined on OC43 spike protein. Panel F: Normalized semi-covariance for generalized mass-charge ratio of human seasonal coronaviral HKU1 (GenBank: MN488637.1) baselined on OC43 spike protein.

Table 3 presents the detailed analysis results for the comparison of OC43 spike protein with the spike proteins from murine coronaviruses, showing that Mass-Charge variances reveal more dependency and trend of each protein sequence evolution [28]. From the analyses, the virulence of MHV strains AB551247.1 and GU593319.1 is similar to the strain FJ884686.1 at ~60%; while the virulence of MHV strain FJ647218.1 (42.45%) is less than strain FJ884686.1.

Fig. 5 and Fig. 6 show the effects of different Z values (binding distance) for different strains of coronaviruses. The valley (binding depth) of the virulence for the different strains of viruses is slightly different based on the previous analysis. For example, in Fig. 5, the Omicron variant has a valley with a similar high magnitude to BA5 and Delta. However, a slightly bigger horizontal Z position value than the Delta variant suggests its lower virulence. Fig. 6 shows the effect of different Z values based on the angle (binding width). The larger the Z value and the higher the angle is, the more contagious the virus. The SARS-CoV-1 shows the lowest angle, while the Omicron variant BA2 and BA5 with the highest Z value is the most infectious variant known.

**Fig. 5.**
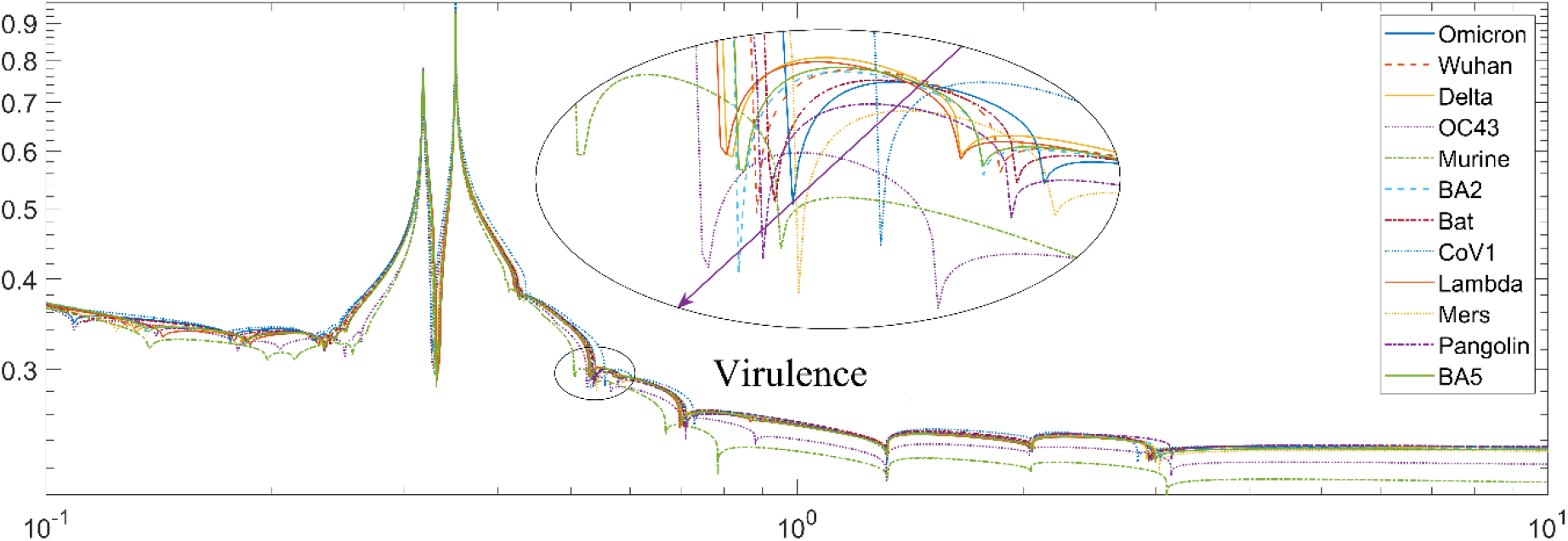
Effects of different distance Z of the Langlands program on viral virulence (amplitude). The curves highlighted in the ovary circle represent the M/Q ratio for optimal distance to interact with the receptors. The arrow direction represents the increase in virulence.

**Fig. 6.**
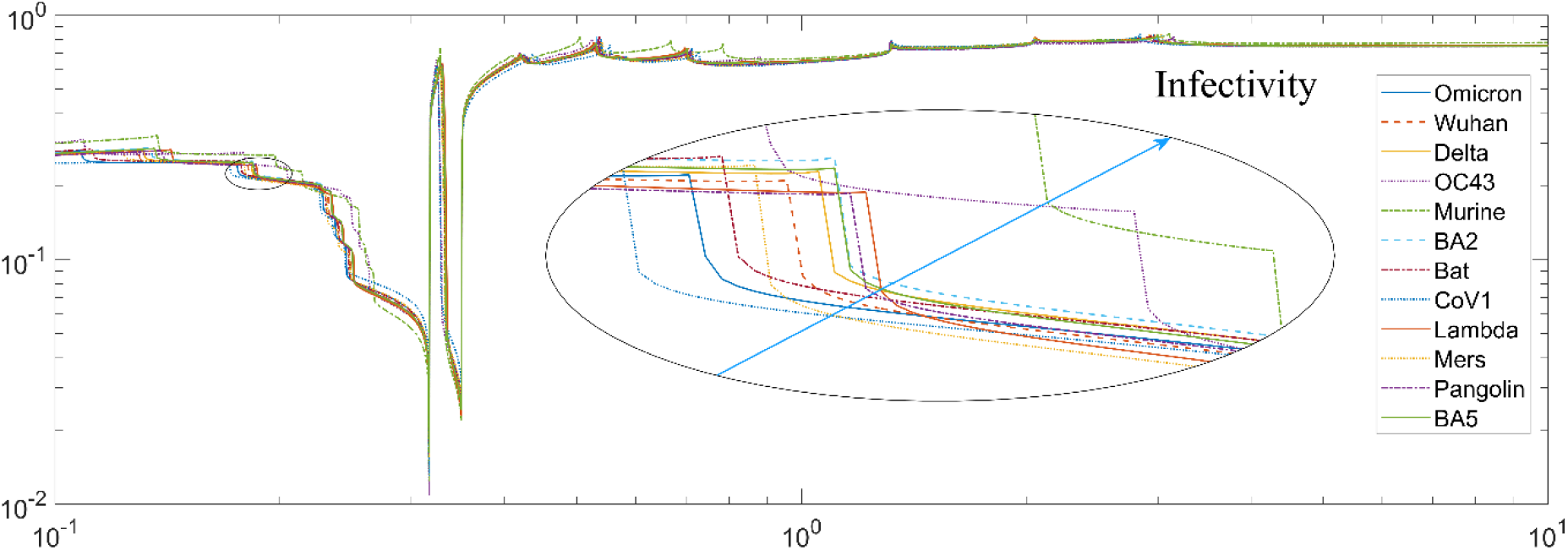
Effects of different distance Z of the Langlands program on viral infectivity (angle). The arrow direction represents the increase in infectivity.

Based on our analysis of the binding coverage for each variant in the past months, we can draw the trend line to demonstrate the evolution of the coronaviruses (Fig. 7). If the binding coverage drops to zero, the viruses may not need further mutations in this region for its infectivity and virulence or the virus may be stabilized relatively. Alternatively, further mutations may not significantly increase the viral infectivity or virulence.

**Figure 7.**
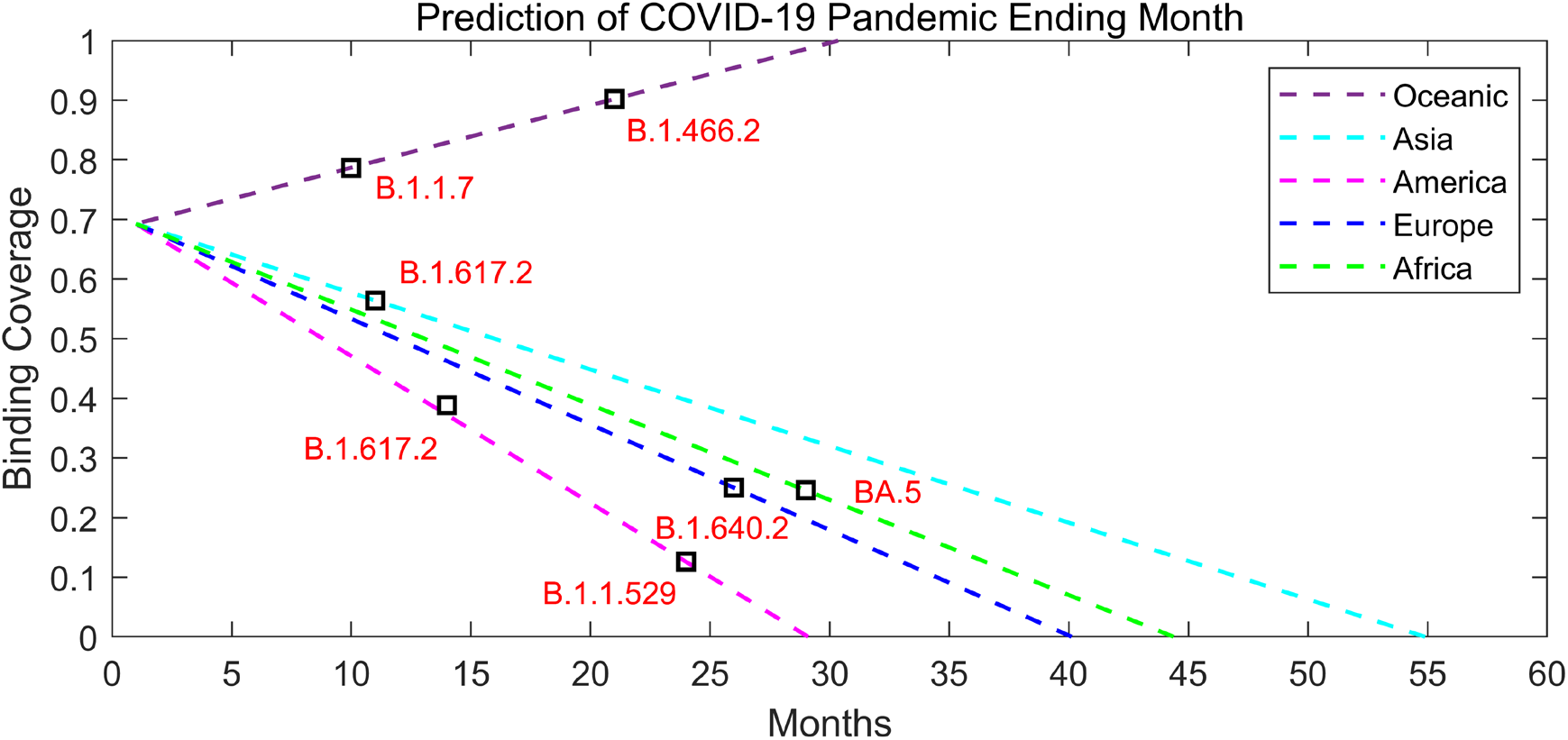
Comparison of different active centers of convergence for SARS-CoV-2 variant spike proteins vs. appearing month and its linear prediction of viral stabilization.

## Conclusion

This study presents the construction of a Mass-Charge covariance for equivalent analysis of coronaviral spike protein fluctuations using an equivalent moment basis with a simple algorithm coded in Excel, and other similar tools can be used too. This novel Mass-Charge model reveals an extra performance index over the traditional model, like infective reproduction rate and virulence estimation. The study also compares the spike proteins from murine coronaviruses to the spike proteins from human common cold seasonal coronaviruses. The results of the Mass-Charge calculation show the differences between animal and human coronaviral spike proteins that traditional covariance definition and calculation may overlook. It also reveals the unique positive and negative charge (mutation) section center positions in the viral spike proteins (where the convergence and divergence sections are located). By using the normalization procedure, the new algorithm removes the self-correlation of each viral spike protein and displays only the cross-correlation between the viral spike proteins. By analyzing various parameters of viral protein sequences using the algorithm, the infectivity and the virulence of the viruses or viral variants may be accurately predicted, particularly when a new virus or a new viral variant is emerging. If the viral infectivity and virulence can be estimated or predicted, it will provide important information for preventative measures or therapeutic preparedness for the diseases caused by the virus and its variants.

It is envisioned that the Mass-Charge model is a promising alternative for the coronaviral spike protein analysis as well as for other human and viral protein analyses. The model will be useful to combine inter-virus and intra-virus characterizations. The simplified Excel calculation is very easy to use, accurate enough, and forward compatible with the traditional Pearson model and calculations. The more complicated Matlab code is good for experienced users to do more deep analyses for viral biology and evolution and for drug development. The example code is available from an Excel file on the GitHub and Matlab servers: (https://github.com/steedhuang/Poincare-Fuchs-KleinAutomorphicFunction-COVID19-Mutations). (https://www.mathworks.com/matlabcentral/fileexchange/106870-calculate-langlands-automorphic-mass-charge-spectrum).

## Acknowledgement

The research work conducted by Dr. W. Zhang was supported by funding from the National Research Council of Canada and by a team grant from the Canadian Institute of Health Research (CIHR) on the Rapid Research Response to COVID-19 Outbreak to Dr. Marc-Andre Langlois at the University of Ottawa.

## Author contributions

TX and SZ conducted data analyses, drafted the manuscript and prepared the figures. JSH developed the algorithm, conceived the idea, wrote the manuscript and prepared the figures. WZ conceived the idea, collected data for coronaviral spike proteins, wrote the manuscript and prepared a figure, and finalized the manuscript.

## Code availability

The example code is available from an Excel file on the GitHub server: (https://github.com/steedhuang/Poincare-Fuchs-KleinAutomorphicFunction-COVID19-Mutations). (https://www.mathworks.com/matlabcentral/fileexchange/106870-calculate-langlands-automorphic-mass-charge-spectrum).

